# Antimicrobial and Antioxidant Activity of Ethanolic Extracts of Charred and Uncharred Skins of Four-Toed Hedgehog, *Atelerix albiventris* (Erinaceidae) used in Traditional Management of Boils in Ghana

**DOI:** 10.1101/2025.04.02.646865

**Authors:** Evans Paul Kwame Ameade, Augustine Mra Yaw Aidoo, Francis Kwaku Dzideh Amankwah, Emmanuel Adom

**Author notes:** Email addresses: Augustine Mra Yaw Aidoo, Francis Kwaku Dzideh Amankwah, Emmanuel Adom. Corresponding author: Evans Paul Kwame Ameade, Department of Pharmacognosy and Herbal Medicine, School of Pharmacy and Pharmaceutical Sciences, University for Development Studies, P.O. Box TL 1350, Tamale.

## Abstract

**Aim:** The charred dry skin of the Four-toed Hedgehog (*Atelerix albiventris*) is used as traditional medicine to treat boils. This study was to validate the antimicrobial activity of ethanolic extract of charred dry skin of *Atelerix albiventris* against microorganisms implicated in boils and to determine the antioxidant activity of these extracts. We also aimed to assess if the ethanolic extract of the uncharred skin possesses comparable bioactivity to the charred sample.

**Methodology:** Well diffusion and broth dilution methods were used for the antimicrobial activity, recording the zone of inhibition, Minimum Inhibitory Concentration and Minimum Bactericidal/Fungicidal concentrations of the extracts and the controls. The antioxidant activity was determined by employing 1,1-diphenyl-2-picrylhydrazyl (DPPH) free radicals scavenging and the Total Antioxidant Capacity (TAC) assays.

**Results:** The test results showed that the charred extracts possess significantly antimicrobial activity against the tested microbes than the uncharred samples. Against the fungus *Candida albicans*, the charred extract recorded a zone of inhibition of 19 mm at 50 mg/ml, while against the main boil-causing bacteria, *Staphylococcus aureus*, the charred extract showed activity even at 25 mg/ml (10.7 mm) with the uncharred extract only becoming bioactive at 250 mg/ml concentration (14.7 mm). The DPPH scavenging activity showed that the IC_50_ is 133.40 µg/ml for ascorbic acid, 210.43 µg/ml for charred and 138.93 µg/ml for the uncharred sample. On the other hand, the total antioxidant capacity of the charred sample was higher with Gallic Acid Equivalent Concentrations of 452.03 GAE mg/g as compared to the uncharred sample with 30.75 GAE mg/g.

**Conclusion:** The results of this study validate the traditional use of the skin of *Atelerix albiventris* for the treatment of boils in Ghana. The significantly higher antimicrobial and antioxidant activity exhibited by the extract of the charred sample over the uncharred also validates the method of preparation traditionally employed.

## INTRODUCTION

Traditional medicine (TM) is a foundational component of global health, encompassing indigenous knowledge, skills, and practices aimed at disease prevention, diagnosis, and treatment, as recognized by the World Health Organization (WHO, 2019). Approximately 80% of populations in developing countries depend heavily on TM as a primary form of healthcare due to limited access to modern facilities (Bruschi et al., 2011; WHO, 2019). Although plant-based remedies are often central to TM, the medicinal use of animals and animal products known as zootherapy remains a crucial, albeit under-researched, aspect of traditional medical systems. Zootherapy is widely practised across continents, from the Americas to Asia and Africa, where animal-based remedies are employed to treat conditions ranging from infections to chronic diseases (Policarpo et al., 2019; Ahmad et al., 2023). Although to a limited extent, zootherapy finds some roles in the healthcare system of some developed countries. Traditional Chinese Medicine (TCM) utilizes animal and parts like deer antlers, tiger bones, antelope, buffalo or rhino horns, deer, testicles and os penis of dog, bear or snake bile and seahorses for treatment such as joint pain, infertility and others (Still, 2003; Yu et al., 2022). Marine-derived therapies, like shark cartilage for joint health and cancer, are available in the U.K., Japan, and other European countries (Mahomoodally et al., 2021). Benítez et al., (2012T) reported some folkloric use of snakeskins to treat colds in some parts of Spain. In many developing countries, zootherapy is deeply rooted in traditional practices and is often integrated into the primary healthcare system. The reliance on traditional medicine, including zootherapy, is particularly prevalent in rural and underserved areas where access to modern medical facilities is limited (Fokunang et al., 2011). For example, in the vicinity of Sambulangan Village in the North Bulagi District of the Banggai Islands Regency in Indonesia, a myriad of therapeutic organisms are utilized, including the monitor lizard, snake, sand crab, earthworm, honey bee, and snakehead fish (Gobana et al., 2019). Also, Latin American countries such as Brazil, Mexico, Bolivia and Puerto Rico use animals within the *Porifera and Cnidarian groups*, species such as *Spongia officinalis* and *Physalia physalia* traditionally to treat ailments like asthma and diarrhoea. Traditional medicines in Africa, commonly sold in both rural and urban markets, use a variety of wild animal and plant species as primary ingredients. Numerous wild animal species, including mammals, reptiles, and birds are utilized for their perceived medicinal and curative properties in diseases. Animal parts such as bones, fat, skin, and intestines are believed to treat various ailments ranging from mental and physical illnesses to antenatal care (Ntiamoa-Baidu, 1992). In South Africa, the pangolin is particularly valued for its medicinal properties by the Lobedu tribe (Maliehe,1993). The Iraqw people in Northern Tanzania and communities in the Assosa Districts of Ethiopia also utilize various wildlife species for sustenance, healing, and ritualistic intentions (Green et al., 2022; Manqele et al., 2023). In Nigeria and Ghana, traditional healers use parts from animals like leopards, elephants, Chameleon, tortoises, Mud-fish (*Clarias sp*.) and pythons for treating a wide range of conditions such as rheumatism, back pain, and convulsions (Adeola, 1992; Ntiamoa-Baidu, 1992).

In recent years, scientific interest has grown in assessing the bioactive compounds within animal-derived substances, driven by the need to validate and standardise traditional remedies (Friant et al., 2022; Daimari & Dutta, 2023). Research into zoochemicals has revealed that many animal tissues contain alkaloids, saponins, polyphenols, and other compounds with potential antioxidant and antimicrobial activities (Villabeto et al., 2019; Thamizharasan & Ravichandran, 2023). The Four-toed hedgehog (*Atelerix albiventris*), a small omnivorous mammal of the family Erinaceidae is traditionally used in Ghana for the treatment of skin ailments including boils. The skin is charred, powdered and applied to the affected part of the skin. There is currently no study to validate this traditional use and preparation method hence this research aims to evaluate the antimicrobial and antioxidant activities of ethanolic extracts from charred and uncharred skins of *Atelerix albiventris*. Through this investigation, we seek to bridge the gap between indigenous knowledge and contemporary science, potentially uncovering new bioactive compounds with applications in integrative medicine.

## MATERIALS AND METHODS

### Study Design

This research employed an experimental laboratory design to determine the antimicrobial activity and to determine the antioxidant properties of the skin of the Four-toed hedgehog (*Atelerix albiventris*) through antioxidant Assays.

### Sample Collection and Identification

The Skin of Four-toed hedgehog (*Atelerix albiventris*) was purchased from the Tamale central market in the northern region of Ghana. The dried skin (figure 1) was identified and authenticated at the Department of Pharmacognosy and Herbal Medicine, School of Pharmacy and Pharmaceutical Sciences, University for Development Studies, Tamale. Figure 1 shows pictures of the dried skin of *Atelerix albiventris*.

**Figure 1:**
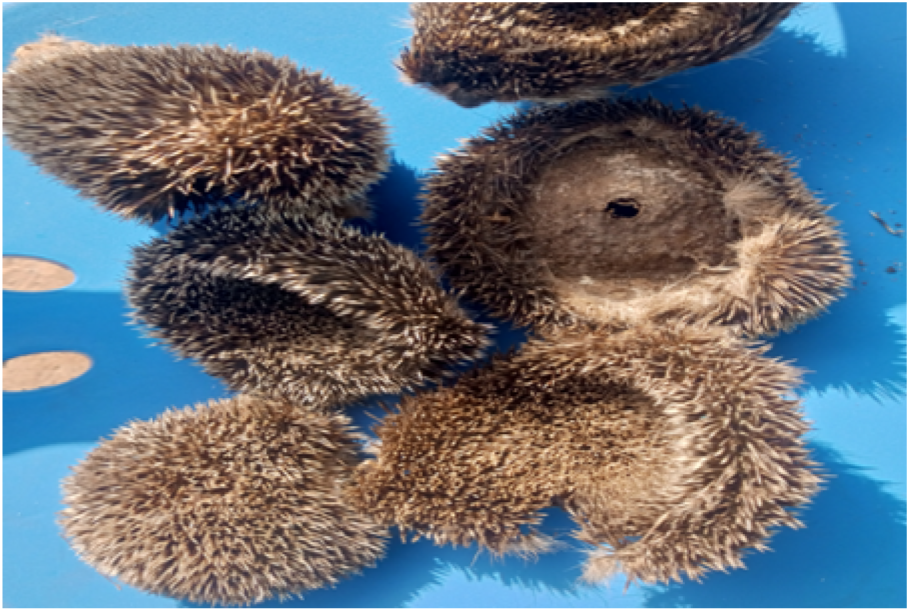
Picture of the dried skins of *Atelerix albiventris*.

### Preparation of Extract

A cold maceration method with few modifications was used to produce the extract. The skins of were further dried in the oven at 40°C for five days. The extracts were prepared from ten dried skins of *Atelerix albiventris*; the two samples from the skins of *Atelerix albiventris* were prepared by charring five dry skins which were then allowed to cool and finally ground into fine powder by mortar and pestle and the other five were also cut and grounded using an electrical blender into skin powder and stored at room temperature. The charring was done by cutting the skins into smaller pieces which were burnt in a metallic bowl placed on a Hotplate stove. The powdered material of the charred and uncharred skin were extracted with 300 ml and 500 ml of 70% ethanol respectively using maceration method for 48 hours with occasional agitation in a stoppered amber bottle. The extracts were filtered using a filter pump and concentrated using a rotary evaporator followed by lyophilization for 24 hours in a freeze dryer (Azwanida, 2015; Zhang et al., 2018). Figures 2 and 3 represent grounded uncharrred dried skin and grounded charred skin of A. albiventris respectively

**Figure 2:**
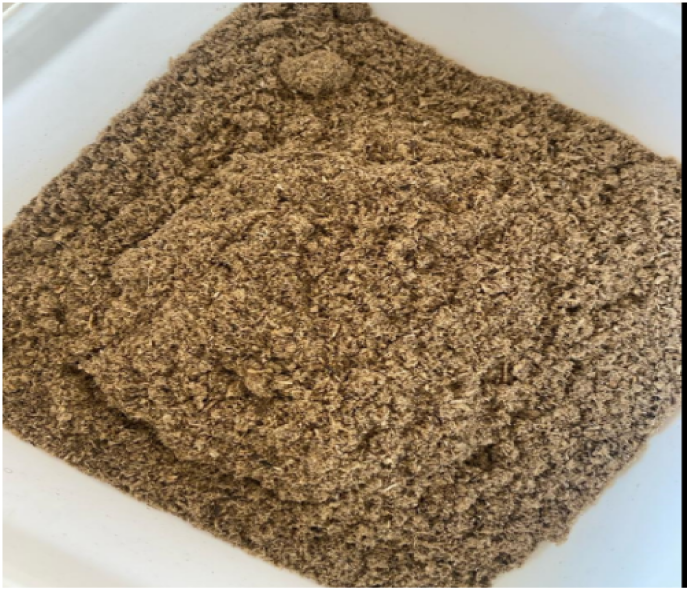
Powdered uncharred sample of *A. albiventris*

**Figure 3:**
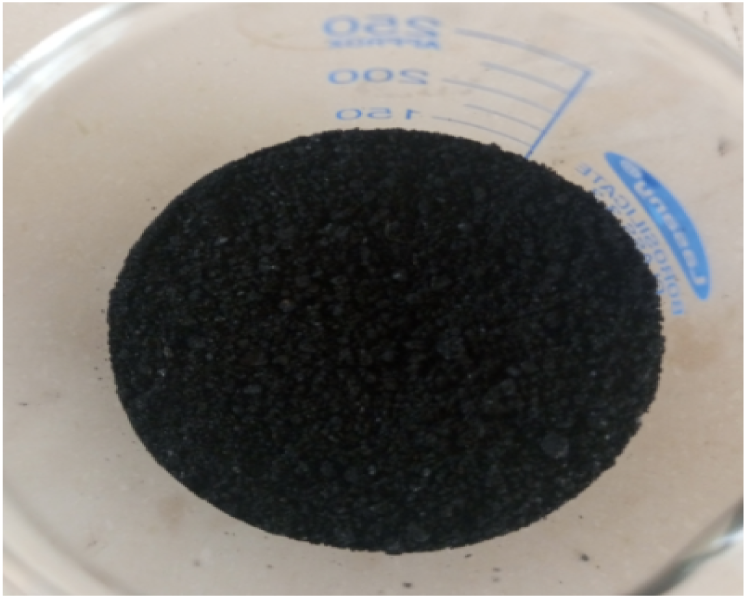
Powdered charred sample of *A. albiventris*

### Calculation of Extraction Yield

The % yield of the charred and uncharred skin extract was calculated using the formula;

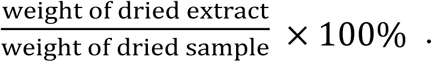

### Test organisms

The bacterial cultures of *Staphylococcus aureus, Streptococcus pyogenes, Pseudomonas aeruginosa, Candida albicans and Methicillin-resistant Staphylococcus aureus* was obtained from the Department of Pharmaceutics, School of Pharmacy and Pharmaceutical Sciences, University for Development Studies, Tamale. All bacterial cultures was be stored in Nutrient agar (NA) slants at 4°C and incubated at 35°C for 18-24 hours. The microbial pathogens was stored and handled in the microbiology laboratory with Biosafety level 2 (BSL-2).

### Antimicrobial activity

All test organisms were sub-cultured in nutrient broth, and the antimicrobial activity of the ethanolic extract from the skin of *Atelerix albiventris* was initially evaluated using an agar well diffusion assay. To prepare the stock solution at a concentration of 400 mg/ml, 4 g of each extract (both charred and uncharred) was dissolved in a 10 ml 3:2 distilled water and DMSO solution. This stock solution was then serially diluted to achieve concentrations of 300 mg/ml, 250 mg/ml 200 mg/ml, 150 mg/ml, 100 mg/ml, 50 mg/ml, 25 mg/ml, and 12.5 mg/ml. The antimicrobial effects of the extracts were tested at these concentrations. For each concentration, 0.2 ml of the extract solution was dispensed into a well.

Nutrient agar was used as the growth medium for both bacteria and fungi. The medium was prepared according to the manufacturer’s instructions, sterilized at 121°C for 15 minutes, and then cooled to 48°C in a water bath. A volume of 100 ml of the prepared nutrient agar was poured into petri dishes containing 100 µL of the test organism, which was swirled to ensure even distribution before being allowed to cool and solidify. Wells with an 8 mm diameter were created using sterilized borers. Each well was filled with 0.2 ml of the extract solution, allowed to diffuse for 1 hour, and then incubated at 37°C for 24 hours. Following incubation, the medium was examined for zones of growth inhibition.

As controls, wells filled with the 3:2 mixture of distilled water and DMSO served as negative controls, while wells containing 30 µg/ml chloramphenicol for bacteria and fluconazole for fungi served as positive controls. The experiment was conducted in triplicate, and the zones of inhibition were measured in millimeters using a transparent ruler (Yusuf et al., 2015). The mean values were then recorded.

### Determination of minimum inhibitory concentration (MIC)

The broth microdilution method was employed to determine the Minimum Inhibitory Concentration (MIC) of the test extracts, adhering to the Clinical and Laboratory Standards Institute (CLSI) document M07-A9, 2012 (CSLI, 2012).

### Materials

The following materials were used for this procedure:

Sterile 96-well microtiter plates, nutrient broth (for bacterial analysis), serial dilutions of the test extracts, standardized inocula of test organisms (0.5 McFarland standard), pipettes and sterile tips and incubator.

### Procedure

#### Preparation of Dilutions

Growth medium (Nutrient Broth) was dispensed into the wells of a 96-well microtiter plate. A starting concentration of the test extract was added to the first column of wells. Two-fold serial dilutions were performed across the rows, creating a decreasing concentration gradient of the extract.

#### Inoculation

A standardized volume of 80 µL of microbial inoculum was added to each well, except the negative control wells.

#### Controls

The following controls were included;

Positive Control: Chloramphenicol 100 mg/ml and Fluconazole 1000 µg/ml.

Negative Control: A well containing growth medium and test organism only was used as control.

#### Incubation

The plates were incubated at the appropriate temperature for the specified duration: 37°C for 24 hours After incubation, the plates were visually examined. The well with the lowest extract concentration that exhibited no visible growth was identified as the Minimum Inhibitory Concentration (MIC).

#### Biological samples disposal

After the execution of the microbiological assays, glassware containing bacterial cultures were all autoclaved at 121° C and 15 psi (1.02 atm). Similarly, the same decontamination and sterilization procedures were followed for all of the glassware used. Waste and used materials would be disposed of properly in waste bags intended for biological samples provided by the Microbiology Laboratory of the University for Development Studies, Tamale.

### Antioxidant Assays

#### Determination of 1,1, dipheny-2-picrylhydrazyl (DPPH) Radical Scavenging Activities

A DPPH radical scavenging assay was conducted using 1,1-diphenyl-2-picrylhydrazyl (DPPH) as outlined in Brand-Williams et al.’s method, with a few adjustments. Five concentrations of plant extracts (0.0625, 0.125, 0.25, 0.5, and 1 mg/ml) were dissolved in analytical-grade methanol. The same concentration range was also prepared for L-ascorbic acid, which served as a standard reference for antioxidant activity. For each sample, 1 ml of the plant extract was placed in a test tube, followed by the addition of 0.5 ml of a 0.3 mM DPPH solution in methanol. The mixture was shaken, then set aside in a dark environment at room temperature for 15 minutes. To ensure accurate measurements, blank solutions were prepared, consisting of 2.5 ml of each plant extract solution mixed with 1 ml of methanol as a baseline. The negative control included 2.5 ml of DPPH solution and 1 ml of methanol, while the positive control used L-ascorbic acid at the same concentrations as the extracts. After the dark incubation period, absorbance was recorded at 517 nm using a spectrophotometer. All tests were conducted in triplicate. The DPPH radical scavenging activity was calculated with the equation provided by Brand-Williams et al.:

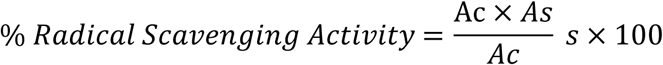

where As is the absorbance of the sample, and Ac is the absorbance of the control. The half maximal inhibitory concentration (IC_50_) of the extracts was computed from a plot of percentage DPPH free radical inhibition versus the extract concentration.

#### Total Antioxidant Capacity (TAC) Assay

The total antioxidant capacity of the ethanolic extracts from both charred and uncharred skin of *Atelerix albiventris* was analyzed using a modified version of Prieto et al.’s method. This technique relies on the reduction of phosphomolybdic acid (Mo (VI)) to a blue phosphomolybdenum (Mo (V)) complex by the test extracts and a reference standard. Gallic acid served as the standard reference, with distilled water used as the blank. A stock solution of Gallic acid at 100 µg/mL was prepared, from which additional concentrations of 50, 25, 12.5, and 6.125 µg/mL were created through serial dilution in sterile distilled water. Separately, a test solution of each extract was prepared at a concentration of 500 µg/mL in sterile distilled water. For each reaction mixture, 5 mL of the test solution was combined with 5 mL of phosphomolybdenum reagent (comprising 0.6 M sulfuric acid, 28 mM sodium phosphate, and 4 mM ammonium molybdate) in a series of test tubes, making a total volume of 10 mL per sample. The samples were then incubated in a 95°C water bath for 90 minutes. After cooling, absorbance readings for each solution were taken in triplicate using a UV-visible spectrophotometer set to 695 nm. This entire procedure was repeated independently to produce three sets of data for thorough analysis. A calibration curve was generated by plotting the absorbance of ascorbic acid solutions against their concentrations using GraphPad software. The absorbance values of the extract solutions were substituted as the dependent variables into the linear equation from the Gallic acid curve. This calculation allowed for the determination of Gallic acid equivalents (GAE), presented as GAE/100 mg/g Gallic acid.

## Results and Analysis

### Extraction Yield of Samples

Table 1 shows the percentage yield of extracts from charred and uncharred dried skin samples and their respective weights. The percentage yield of extracts obtained from both charred and uncharred dried skin samples varied. For the charred sample, the weight after drying was 46.81 g and the extract obtained from this sample weighed 9.41 g. This resulted in a percentage yield of 20.10%. In contrast, the uncharred sample had a higher weight after drying, recorded at 76.67 g, but yielded a lower extract weight of 6.14 g, resulting in a per cent yield of only 8.00%.

**Table 1.** Percentage Yields of Extracts.

| Sample | Weight of Dried Sample(g) | Weight of Extract(g) | Percentage Yield(%) |
| --- | --- | --- | --- |
| Charred | 46.81 | 9.41 | 20.10 |
| Uncharred | 76.67 | 6.14 | 8.00 |

### Antimicrobial Activity

#### Zone of inhibition

Table 2 presents the mean zone of inhibition for the charred and uncharred samples. Both charred and uncharred samples exhibited inhibition at higher concentrations (250-300 mg/ml) against the tested organisms like *Staph. aureus* and *P. aeruginosa*. Inhibition varied across concentrations for both charred and uncharred samples. For *Staph. aureus*, the charred sample showed inhibition starting at 25 mg/ml with a mean zone of 10.67 mm, increasing to 19.67 mm at 300 mg/ml. The uncharred sample began inhibition at 250 mg/ml, with a zone of 14.67 mm at 250 mg/ml and 19mm at 300 mg/ml. For *Strep. pyogenes*, the charred sample started inhibiting at 25 mg/ml with a zone of 9.33 mm, while the uncharred sample showed inhibition at 250 mg/ml with 14.33 mm and 15.33mm at 300 mg/ml. For *P. aeruginosa*, the charred sample inhibited at 25 mg/ml with a zone of 11.67 mm, whereas the uncharred sample started at 150 mg/ml. Both samples showed inhibition at 250 mg/ml and 300 mg/ml. Against MRSA, the charred sample showed inhibition at 25g/ml with 11.00mm, while the uncharred sample begun at 150g/ml. For *C. albicans*, the charred sample shows inhibition at 50 mg/ml with 19.00 mm, and the uncharred sample starts at 150 mg/ml with 14.67 mm. Significant differences (p-value < 0.001) were observed between the charred and uncharred extracts at 25 mg/ml, 50 mg/ml, 100 mg/ml, and 150 mg/ml against *Staphylococcus aureus*. However, the differences were not statistically significant at higher concentrations of 200 mg/ml and 300 mg/ml (p-value = 0.116). Similarly, no significant differences in zones of inhibition were observed in MRSA at concentrations of 200 mg/ml, 250 mg/ml, and 300 mg/ml, with p-values of 0.561, 0.643, and 1.00 respectively. Pictures of the agar well diffusion test are shown in Figure 4.

**Table 2:** Mean Zone of Inhibition of Charred and Uncharred Sample.

| Test Organism | Concentration(g/ml) | Mean Zone of Inhibition± SD(mm) |  | P-value |
| --- | --- | --- | --- | --- |
|  |  | Charred | Uncharred |  |
| *Staph. aureus* | 12.5 | 0.0±0.0 | 0.0±0.0 | N/A |
|  | 25 | 10.7±1.2 | 0.0±0.0 | <0.001 |
|  | 50 | 13.0±1.0 | 0.0±0.0 | <0.001 |
|  | 100 | 13.7±1.5 | 0.0±0.0 | <0.001 |
|  | 150 | 13.7±0.6 | 0.0±0.0 | <0.001 |
|  | 200 | 14.0±0.0 | 0.0±0.0 | 0.116 |
|  | 250 | 16.0±1.0 | 14.7±0.6 | 0.012 |
|  | 300 | 19.7±0.6 | 19.0±0.0 | 0.116 |
|  | Positive Control | 21.7±0.6 | 22.3±0.6 | 0.230 |
| *Strep. pyogenes* | 12.5 | 0.0±0.0 | 0.0±0.0 | N/A |
|  | 25 | 9.3±0.6 | 0.0±0.0 | <0.001 |
|  | 50 | 10.0±1.0 | 0.0±0.0 | <0.001 |
|  | 100 | 12.0±1.7 | 0.0±0.0 | <0.001 |
|  | 150 | 12.0±1.0 | 0.0±0.0 | <0.001 |
|  | 200 | 13.3±0.6 | 0.0±0.0 | <0.001 |
|  | 250 | 16.3±0.6 | 14.3±0.6 | 0.013 |
|  | 300 | 19.3±1.5 | 15.3±1.2 | 0.022 |
|  | Positive Control | 21.0±1.0 | 22.7±1.0 | 0.067 |
| *P. aeruginosa* | 12.5 | 0.0±0.0 | 0.0±0.0 | N/A |
|  | 25 | 11.7±1.2 | 0.0±0.0 | <0.001 |
|  | 50 | 14.3±2.1 | 0.0±0.0 | <0.001 |
|  | 100 | 16.3±0.6 | 0.0±0.0 | <0.001 |
|  | 150 | 19.0±1.7 | 12.3±1.6 | 0.007 |
|  | 200 | 20.0±1.0 | 16.3±1.5 | 0.025 |
|  | 250 | 20.0±2.0 | 20.0±0.0 | 1.00 |
|  | 300 | 22.3±1.5 | 20.7±0.6 | 0.152 |
|  | Positive Control | 20.3±1.2 | 23.0±1.0 | 0.039 |
| *MRSA* | 12.5 | 0.0±0.0 | 0.0±0.0 | N/A |
|  | 25 | 11.0±0.0 | 0.0±0.0 | <0.001 |
|  | 50 | 12.3±0.6 | 0.0±0.0 | <0.001 |
|  | 100 | 14.3±0.6 | 0.0±0.0 | <0.001 |
|  | 150 | 15.7±0.6 | 11.7±0.6 | 0.001 |
|  | 200 | 17.0±1.0 | 16.3±1.5 | 0.561 |
|  | 250 | 20.3±1.2 | 20.0±0.0 | 0.643 |
|  | 300 | 20.7±1.2 | 20.7±0.8 | 1.00 |
|  | Positive Control | 20.0±1.0 | 19.0±1.0 | 0.288 |
| *C. albicans* | 12.5 | 0.0±0.0 | 0.0±0.0 | N/A |
|  | 25 | 0.0±0.0 | 0.0±0.0 | N/A |
|  | 50 | 19.0±1.0 | 0.0±0.0 | <0.001 |
|  | 100 | 20.7±0.6 | 0.0±0.0 | <0.001 |
|  | 150 | 16.3±0.6 | 14.7±0.6 | 0.024 |
|  | 200 | 17.3±2.5 | 15.0±1.0 | 0.210 |
|  | 250 | 18.0±1.0 | 15.3±1.5 | 0.065 |
|  | 300 | 17.7±1.5 | 13.0±1.0 | 0.011 |
|  | Positive Control | 28.3±2.1 | 26.3±1.5 | 0.251 |

\*N/A = Not applicable. SD = Standard deviation. Concentration of the positive control for the bacteria was 30 mg/ml but that of the fungus was 1000 µg/ml

**Figure 4:**
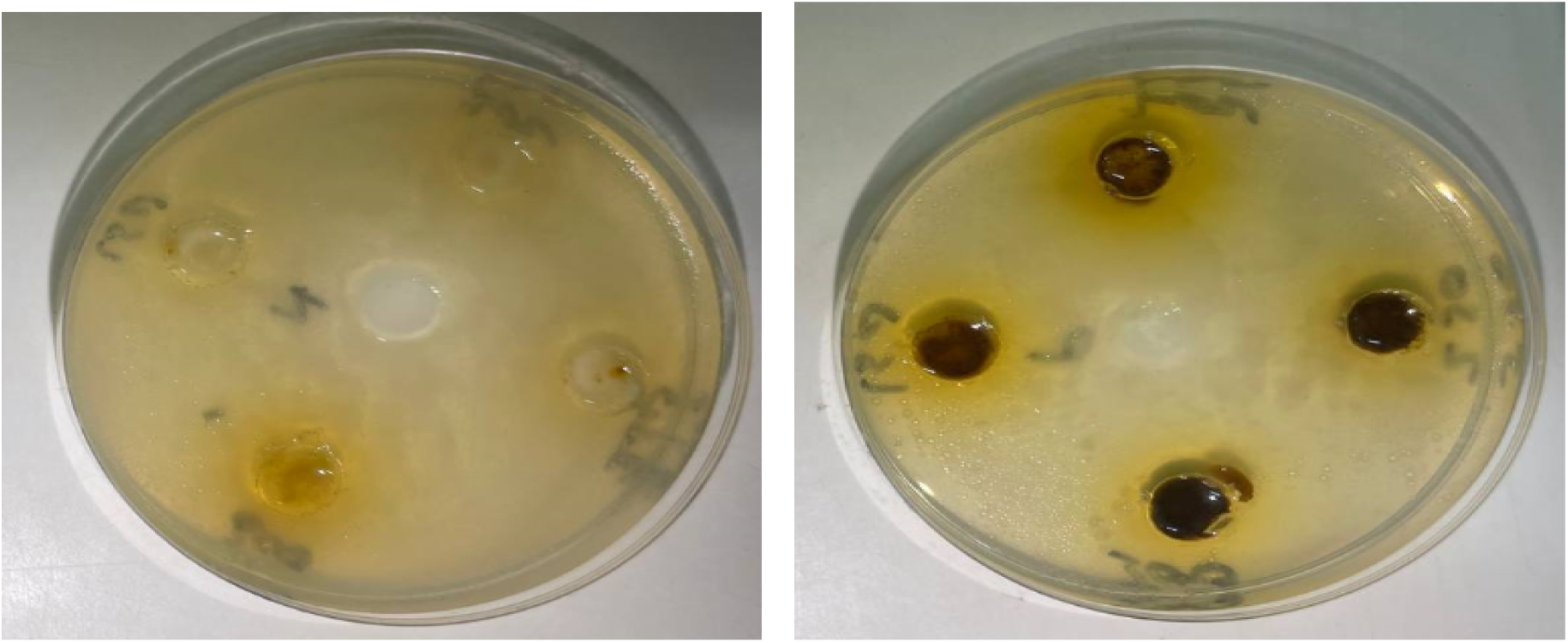
Pictures of Zone of Inhibition from agar well diffusion.

### Minimum Inhibitory Concentration (MIC) of Charred and Uncharred Extracts

The microbial growth was effectively inhibited by both the charred and uncharred samples but at different concentrations. Table 3 shows the MIC values for both the charred and uncharred samples. In both control cases, the inhibitory action at low concentrations is constant. The charred sample was generally associated with lower MIC values (1.562 mg/ml for *Staph. aureus* and *C. albicans*) compared to the uncharred sample, which displayed much higher MIC values (25 mg/ml for *Staph. aureus* and *C. albicans*). Against *Streptococcus pyogenes*, the charred inhibited at 3.343 mg/ml while the uncharred inhibited at 12.5 mg/ml. The charred sample gave MICs of 3.125 mg/ml and 4.688mg/ml and the uncharred 9.375 mg/ml and 12.5 mg/ml against *P*. *aeruginosa* and *MRSA* respectively.

**Table 3:**
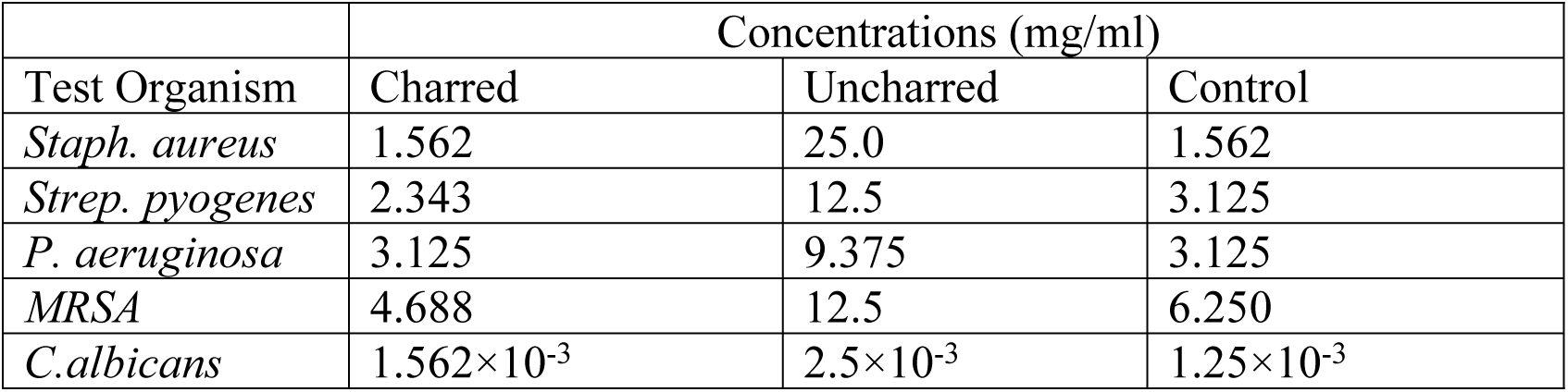
Minimum Inhibitory Concentration (MIC) Of Charred and Uncharred Samples.

| Test Organism | Concentrations (mg/ml) |  |  |
| --- | --- | --- | --- |
|  | Charred | Uncharred | Control |
| *Staph. aureus* | 1.562 | 25.0 | 1.562 |
| *Strep. pyogenes* | 2.343 | 12.5 | 3.125 |
| *P. aeruginosa* | 3.125 | 9.375 | 3.125 |
| *MRSA* | 4.688 | 12.5 | 6.250 |
| *C.albicans* | $1.562 \times 10^{-3}$ | $2.5 \times 10^{-3}$ | $1.25 \times 10^{-3}$ |

### Minimum Bactericidal/Fungicidal Concentration (MBC/MFC)

The charred and uncharred samples showed bactericidal/fungicidal activity. Table 4 details the MIC/MFC of both Charred and Uncharred Samples. They are both capable of complete microbial death at higher concentrations (50-100 mg/ml for most microbes). The charred sample shows lower MBC values (50 mg/ml for *Staph. aureus*, *P. aeruginosa*, MRSA, and *C. albicans*), while the uncharred sample consistently requires 100 mg/ml for complete microbial death.

**Table 4:** Minimum Bactericidal/Fungicidal Concentration (MIC/MFC) Of Charred and Uncharred Samples.

| Test Organism | Concentrations (mg/ml) |  |  |
| --- | --- | --- | --- |
|  | Charred | Uncharred | Control |
| *Staph. aureus* | 50.0±0.0 | 100.0±0.0 | 100.0±0.0 |
| *Strep. pyogenes* | 25.0±0.0 | 100.0±0.0 | 100.0±0.0 |
| *P. aeruginosa* | 50.0±0.0 | 100.0±0.0 | 100.0±0.0 |
| *MRSA* | 50.0±0.0 | 100.0±0.0 | 100.0±0.0 |
| *C.albicans* | 50.0±0.0 | 100.0±0.0 | 1.0×10<sup>-3</sup> ±0.0 |

### Antioxidant Activity

The antioxidant activities of the samples were studied by DPPH scavenging activity. The IC_50_ values of ascorbic acid, uncharred dry skin, and charred dry skin are presented in Table 5, showing that the IC_50_ is 133.40 µg/ml for ascorbic acid. The IC_50_ of the uncharred sample was 138.93 µg/ml. On the other hand, the antioxidant capacity of the charred sample was far lower, with an IC_50_ of 210.43 µg/ml, showing that charring reduced the radical scavenging activity of the sample (Figure 5).

**Table 5:** DPPH scavenging activity.

| Sample | IC<sub>50</sub>µg/ml |
| --- | --- |
| Ascorbic acid | 133.40±10.49 |
| Uncharred | 138.93±28.13 |
| Charred | 210.43±32.63 |

**Figure 5:**
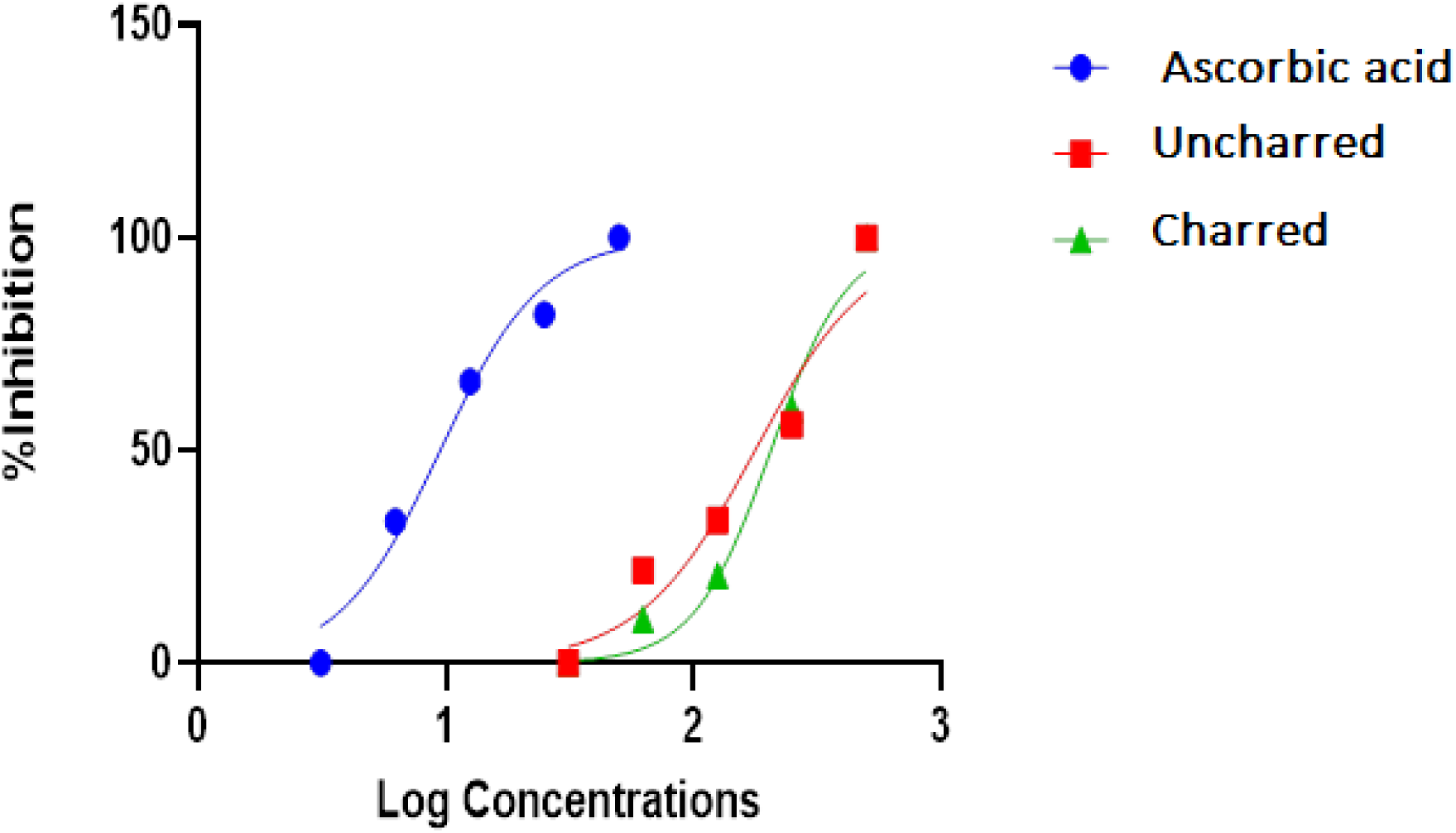
Comparative DPPH radical scavenging activity of the samples with Ascorbic Acid.

Table 6 depicts the Gallic Acid Equivalent (GAE) concentration of charred and uncharred samples The GAE concentration was higher in the charred sample, at 452.03 mg/g ± 0.003, as opposed to a rather low 30.75 mg/g ± 0.05 obtained in the uncharred samples. This sharp contrast reflects that gallic acid concentration has been increased in the charred sample. Figures 6 and 7 represent the plot between absorbance and gallic acid concentration in µg/ml for uncharred and charred samples, respectively. The Figures show the absorbance response of a range of concentrations of gallic acid, indicating a positive linear trend in both cases.

**Table 6:** Gallic Acid Equivalent Concentrations.

| Sample | GAE mg/g |
| --- | --- |
| Charred | 452.03±0.003 |
| Uncharred | 30.75±0.05 |

**Figure 6:**
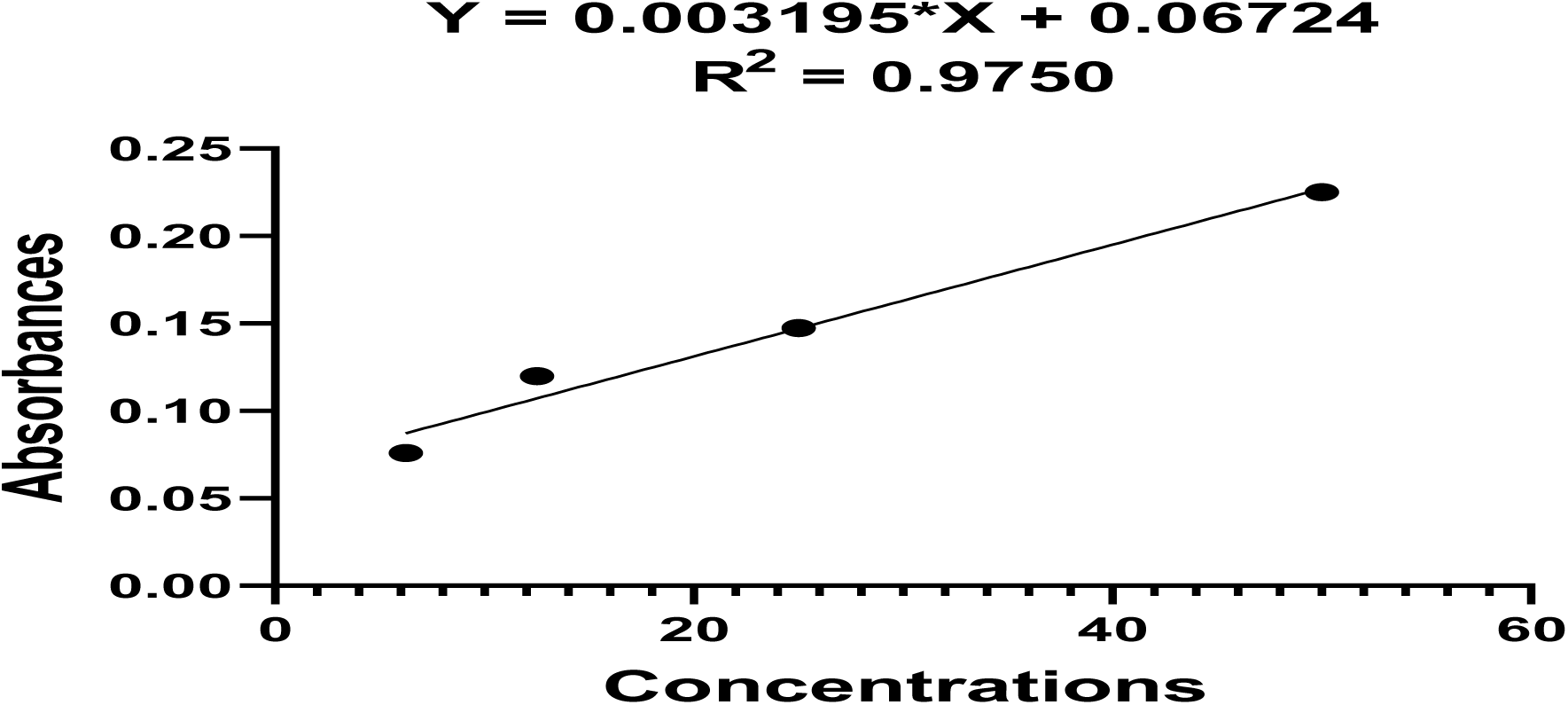
Graph of Absorbance of Gallic Acid Against Concentration (µg/ml) in extract from the uncharred sample

**Figure 7:**
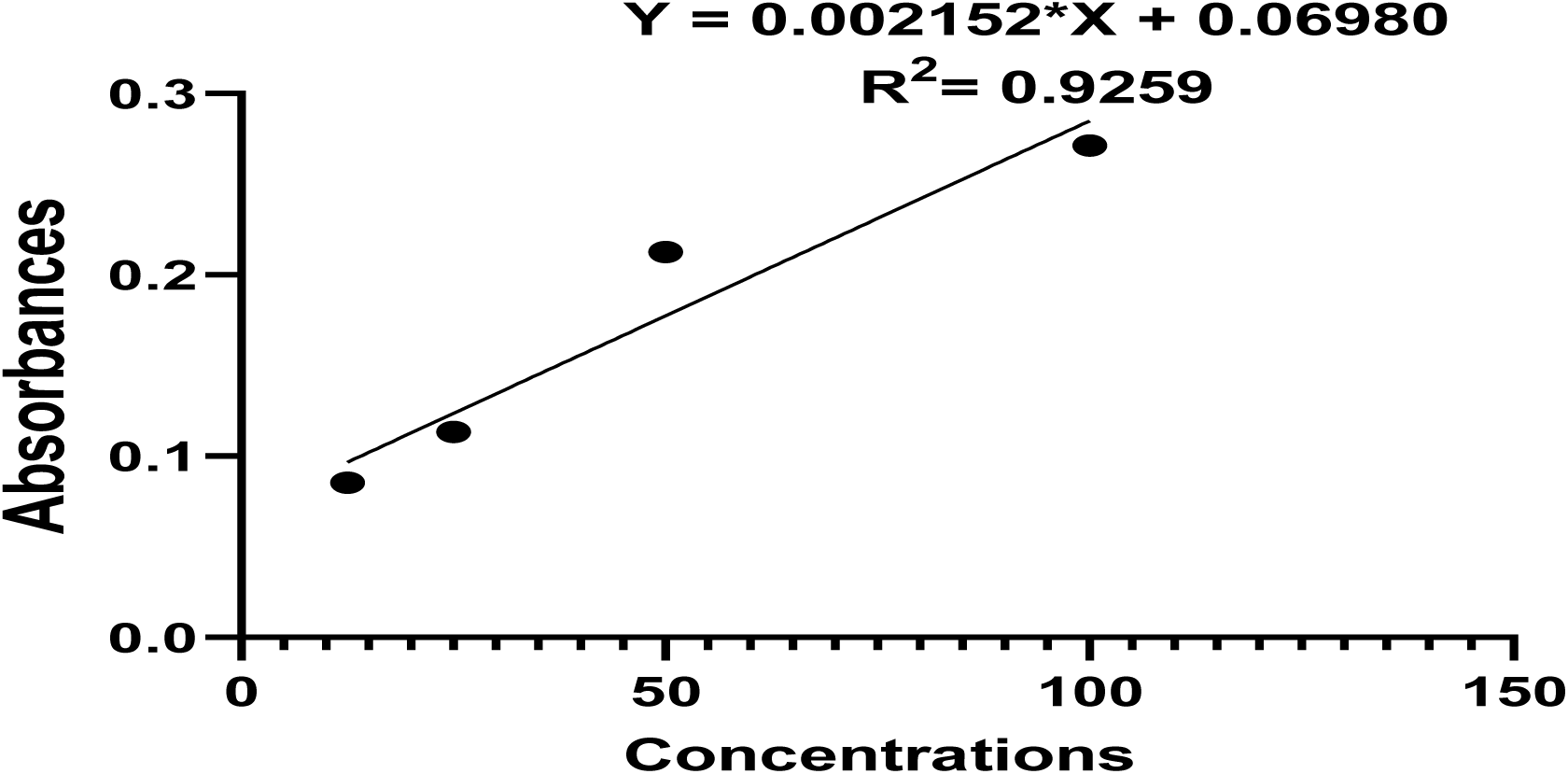
Graph of Absorbance of Gallic Acid Against Concentration (µg/ml) in extract from the charred sample.

## DISCUSSIONS

### Extraction Yield

The charred sample yielded 20.10%, compared to an 8.00% yield from the uncharred sample, indicating that the charring process significantly boosts extraction yield. This increase may be due to structural and chemical changes in the skin caused by charring, making it easier to extract components. This effect is similar to findings by Sebestyén et al. (2019) on tannins, which revealed that high temperatures can cause molecular breakdown or polymerization, altering both structure and properties. Additionally, elevated temperatures are known to improve the solubility of tannins, which can enhance extraction efficiency. Techniques like pressurized hot water extraction, for instance, have shown significantly higher yields of hydrolysable tannins compared to traditional methods. Ubeed et al. (2022) also pointed out that both temperature and extraction duration can influence the efficiency of extracting phytochemicals from plant materials, with higher temperatures and longer extraction times typically yielding better results. Even though the uncharred sample had a greater dry weight (76.67g) compared to the charred sample (46.81g), its extraction efficiency was lower.

### Antimicrobial Activity

The charred sample generally exhibited superior activity at lower concentrations compared to the uncharred sample in the antimicrobial testing. This was evidenced by lower Minimum Inhibitory Concentration (MIC) and Minimum Bactericidal/Fungicidal Concentration (MBC) values, along with larger inhibition zones (ZOI) at reduced concentrations. Charring may have caused the break down of some heat-sensitive compounds in the skin, while enhancing more heat-stable compounds that may possess antimicrobial activity. Thus, charring appears to improve overall antimicrobial effectiveness by removing less stable, less potent compounds and concentrating or producing more effective antimicrobial agents. The differences in antimicrobial capabilities of the two samples could be linked differences in compounds. Overall, the charred sample showed greater antimicrobial efficacy across different organisms and concentrations, as demonstrated by lower MIC and MBC values and larger inhibition zones at reduced concentrations.

When comparing the antimicrobial potential of the charred *Atelerix albiventris* skin extract with the crude methanolic extract of *Perna viridis* (Paduhilao et al., 2022) and the ground beetle extract (Abubakar, 2021), it is evident that various animal extracts display antimicrobial activity. However, their effectiveness against bacteria, such as *Staphylococcus aureus*, varies with concentration. For instance, the charred *Atelerix albiventris* extract showed an average inhibition zone of 19.7 ± 0.6 mm against *S. aureus* at 300 mg/mL, while the *Perna viridis* methanolic extract achieved a similar inhibition zone (19.0 ± 0.9 mm) at a lower concentration of 100 mg/mL in Paduhilao et al.’s study (2022). Against Methicillin-resistant *Staphylococcus aureus* (MRSA), the *Atelerix albiventris* extract recorded an inhibition zone of 20.7 ± 1.2 mm at 300 mg/mL, whereas the methanolic extract from the ground beetle extract reported by Yahaya et al., (2019) caused a smaller inhibition zone of 9.7 ± 0.6 mm at 10 mg/mL. These differences show that although animal extracts can effectively inhibit certain bacteria, their potency differs based on the animal extract, the solvent used for the extraction and concentrations of the test samples.

### Antioxidant Activity

The DPPH radical scavenging and Total Antioxidant Capacity (TAC) assays showed differences in antioxidant activities between the charred and uncharred samples, likely due to variations in their chemical composition. The DPPH assay revealed a lower IC_50_ for the uncharred sample (138.93 µg/mL) than the charred sample (210.43 µg/mL), suggesting that the uncharred sample had stronger antioxidant activity. Both samples were compared with ascorbic acid as a control, which had the lowest IC_50_ value of 133.40 µg/mL (Jumina et al., 2019). In contrast, the TAC showed that while the charred samples had a GAE value of 452.03 mg/g, the uncharred samples measured at only 30.75 mg/g. This implies that charring enhances total antioxidant capacity, despite reducing DPPH scavenging activity. The difference in antioxidant activity may stem from the distinct compounds present in each sample. Charring might degrade certain heat-sensitive antioxidants, which could explain the high TAC but lower DPPH scavenging activity.

## CONCLUSION

In comparing the antimicrobial potency and antioxidant effects of charred versus uncharred samples, it was observed that the charred skin exhibited better antimicrobial effects, particularly against *Staphylococcus aureus*, *Pseudomonas aeruginosa*, and *Candida albicans*. However, the Uncharred sample exhibited a better DPPH radical scavenging activity but lower Total Antioxidant Capacity compared to the charred sample. This study successfully confirmed the antimicrobial and antioxidant effects of the ethanolic extract from the dry skin of *Atelerix albiventris*, especially the charred sample supporting its traditional use and preparation method in Ghanaian traditional medicine.

## ABBREVIATIONS

MRSA: Methicillin Resistant *staphylococcus aureus*
W.H.O: World Health Organization
DPPH: 2,2-diphenyl-1-picrylhydrazyl
DMSO: Dimethyl sulfoxide
MIC: Minimum Inhibitory Concentration
MIC: Minimum Bactericidal Concentration
MFC: Minimum Fungicidal Concentration
ZOI: Zone of Inhibition
TAC: Total Antioxidant Capacity

